# Impact of HIV infection and integrase strand transfer inhibitors-based treatment on gut virome

**DOI:** 10.1101/2022.04.14.488276

**Authors:** Pablo Villoslada-Blanco, Patricia Pérez-Matute, María Íñiguez, Emma Recio-Fernández, Jansen Daan, Lander De Coninck, Lila Close, Pilar Blanco-Navarrete, Luis Metola, Valvanera Ibarra, Jorge Alba, Jelle Matthijnssens, José A Oteo

**Author notes:** Corresponding author: Patricia Pérez-Matute, Infectious Diseases, Microbiota and Metabolism Unit. Infectious Diseases Department, Center for Biomedical Research of La Rioja (CIBIR), Logroño, La Rioja, Spain. C/Piqueras 98, CIBIR building, third floor, 26006 Logroño (La Rioja).

## Abstract

Viruses are the most abundant components of the microbiome in human beings with a significant impact on health and disease. However, the impact of human immunodeficiency virus (HIV) infection on gut virome has been scarcely analyzed. On the other hand, several studies suggested that not all antiretrovirals for treating HIV infection exert similar effects on the gut bacteriome, being the integrase strand transfers inhibitors (INSTIs) —first-choice treatment of naive HIV-infected patients nowadays— those associated with a healthier gut. Thus, the aim of this study was to evaluate the effects of HIV infection and INSTIs in first line of treatment on gut virome composition. To accomplish this objective, 26 non-HIV-infected volunteers, 15 naive HIV-infected patients and 15 INSTIs-treated HIV-infected patients were recruited and gut virome composition was analysed using shotgun sequencing. The results showed that bacteriophages are the most abundant and diverse viruses in the gut independent from the HIV-status and the use of treatment. HIV infection was accompanied by a decrease in phage richness which was reverted after INSTIs-based treatment (p<0.01 naive *vs*. control *Richness* index and p<0.05 naive *vs*. control *Fisher’s alpha* index). β-diversity of phages revealed that samples from HIV-infected samples clustered separately from those belonging to the control group (padj<0.01 naive *vs*. control and padj<0.05 INSTIs *vs*. control). However, it is worth mentioning that samples coming from INSTIs-treated patients were more grouped than those from naive patients. Differential abundant analysis of phages showed an increase of *Caudoviricetes* class in the naive group compared to control the group (padj<0.05) and a decrease of *Malgrandaviricetes* class in the INSTIs-treated group compared to the control group (padj<0.001). Besides, it was observed that INSTIs-based treatment was not able to reverse the increase of lysogenic phages associated with HIV infection (p<0.05 *vs*. control) or to modify the decrease observed on the relative abundance of Proteobacteria-infecting phages (p<0.05 *vs*. control). To sum up, our study describes for the first time the impact of HIV and INSTIs on gut virome and demonstrates that INSTIs-based treatments are able to partially restore gut dysbiosis not only at bacterial but also at viral level, which opens several opportunities for new studies focused on microbiota-based therapies.

**Author summary:** The impact of human immunodeficiency virus (HIV) infection and the effects of integrase strand transfer inhibitors (INSTIs)-based treatments —first-choice treatment of naive HIV-infected patients nowadays— on gut virome are unknown. In this study, we have confirmed that phages are the most abundant viral component of the human gut virome. Besides, we have described for the first time that INSTIs-based treatments are able to partially restore gut dysbiosis induced by HIV infection not only at bacteria but also at viral level. This fact opens new opportunities for future studies and approaches focused on microbiota-based therapies in the context of HIV infection and treatment.

## Introduction

Human immunodeficiency virus (HIV) infection is considered a chronic disease related to a set of structural and functionality changes on the gut epithelial barrier, immunological shifts and modifications in the composition and functionality of gut microbiota (GM) that are not completely restored despite the use of antiretroviral treatments (ARTs) [1,2]. Several studies have analysed the effects of HIV infection and different ARTs on the bacterial component of GM. However, GM consists not only of bacteria but also of viruses, archaea, fungi, and other eukaryotic organisms. In this context, viruses are the most abundant components of the microbiome in human beings with a significant impact on health [3]. In fact, several studies have revealed an altered gut viral composition in different pathological conditions such as cancer [4], type I diabetes [5] and inflammatory bowel disease (IBD) [6]. Bacteriophages, which are the most abundant viruses in the human intestine [7], can also perturb the bacterial community to indirectly influence gut health and interact with the human immune system [8,9]. However, little else is known about the role of bacteriophages and eukaryotic viruses in other human diseases such as HIV infection. Most of the studies assessing the effects of HIV infection on the virome are focused on plasma, saliva, and cervix communities. There is only one study evaluating the effects of HIV infection on the gut virome but it is solely focused on DNA viruses [10], while the importance and impact of RNA viruses cannot be discarded. Besides, it is important to note that the human gut contains 10^9^-10^12^ viral particles [11,12]. On the other hand, we do not know how HIV infection and ARTs act on the gut virome.

Several studies suggest that not all ARTs exert similar effects on the gut bacteriome. In fact, a previous work from our group [13] demonstrated that ARTs based on INSTIs were associated with levels of systemic inflammation, soluble CD14 (sCD14) plasma levels, and microbial diversity similar to those observed in uninfected controls, suggesting a healthier gut and potentially fewer HIV-related complications. In this context, and to our knowledge, there are no studies focused on the effects of the actual ART regimens based on INSTIs, key component of ART in the treatment of naive HIV-infected patients [14–16], on gut virome and its physiological relevance in terms of health. Thus, the objective of this work was to analyse the associations between HIV infection and INSTIs-based therapies in first line of treatment on gut virome composition (both DNA and RNA viruses).

## Results

### Clinical and demographical characteristics of participants

**Table 1** shows the main characteristics of the recruited population. INSTIs-treated patients showed nadir CD4 counts of 526.53 ± 56.30 cells/μl. The average time under treatment was 33.27 ± 5.04 months. Viral load of naive patients was 622,252.9 ± 341,039.3 copies/ml, whereas INSTIs-treated patients showed indetectable viral load (<20 copies/ml), as expected. Statistically significant differences were observed between the naive group and INSTIs-treated patients in terms of CD4 levels (p<0.01) and CD4/CD8 ratio, both higher in the INSTIs-treated group. Statistically significant differences were observed between the control and the naive group in terms of gender (p<0.01), age (p<0.05), systolic blood pressure (p<0.05), diastolic blood pressure (p<0.05) and smoking habits (p<0.05). Thus, males were less represented in the control group (34,62%) in contrast to HIV-infected patients (80% and 86.67% in naive and INSTIs-treated patients, respectively). Mean age of the control group was higher than that observed in the naive group. However, these differences disappeared when comparing the controls against the INSTIs-treated group. No statistical differences on age were observed among the naïve and INSTIs-treated group. Both systolic and diastolic blood pressure were higher in the naive group compared to the control group. No differences were observed when the controls were compared to INSTIs-treated patients and among both HIV-infected groups. Of note, none of the naïve HIV-infected patients referred hypertension. Thus, these differences on blood pressure could be explained as “white coat hypertension” generated in those patients that have received the news of being HIV-positive. Smoking habits were also higher in the naive group and in INSTIs-treated group compared to the control group (11.54% *vs*. 46.67% and 66.67%, respectively). Moreover, no differences were observed neither in the mode of transmission, nor in AIDS events (only one naive patient suffered from it), nor in the coinfection with virus C or B among both HIV groups (only two INSTIs-treated patients presented confection with virus C and a grade of fibrosis of F0/F1).

**Table 1.** Characteristics of healthy uninfected controls and HIV-infected patients (naive and under INSTIs-based treatment).

|  | Control | Naive | INSTIs-treated | p value |
| --- | --- | --- | --- | --- |
| **Number of patients** | 26 | 15 | 15 | - |
| **Gender (men)** | 9/26 (34.62%) | 13/15 (86.67%) ** | 12/15 (80%) ** | **0.002** |
| **Age (years)** | 43.58±2.31 | 33.87±2.85 * | 43.67±3.39 | **0.033** |
| **BMI (kg/m<sup>2</sup>)** | 24.30±0.69 | 23.23±1.05 | 23.51±0.85 | 0.616 |
| **Waist circumference (cm)** | 85.35±2.55 | 83.83±2.86 | 85.13±2.15 | 0.916 |
| **Systolic blood pressure (mm/Hg)** | 120.58±2.76 | 135.67±5.67 * | 129.73±6.02 | **0.050** |
| **Diastolic blood pressure (mm/Hg)** | 72.19±1.94 | 81.87±3.31 * | 77.8±3.24 | **0.035** |
| **Alcohol active** | 3/26 (11.54%) | 0/15 (0%) | 1/15 (6.67%) | 0.578 |
| **Smoking active** | 3/26 (11.54%) | 7/15 (46.67%) * | 10/15 (66.67%) *** | **0.001** |
| **CD4 (cells/μl)** | - | 464.07±76.46 | 850.53±101.68 | **0.006** |
| **CD4/CD8 ratio** | - | 0.53±0.13 | 0.84±0.10 | **0.027** |
| **Mode of transmission** | - | HS: 6/15 (40%) | HS: 7/15 (46.67%) | 0.716 |
|  | - | MSM: 9/15 (60%) | MSM: 7/15 (46.67%) |  |
|  | - | Parenteral: 0/15 (0%) | Parenteral: 1/15 (6.66%) |  |
| **AIDS** | - | 1/15 (6.67%) | 0/15 (0%) | 1 |
| **Coinfection with virus C** | - | 0/15 (0%) | 2/15 (13.33%) | 0.483 |
| **Coinfection with virus B** | - | 0/15 (0%) | 0/15 (0%) | 1 |

Qualitative variables are represented in percentage while quantitative variables are represented as mean ± standard error mean. P value refers to the comparison between two (naive vs. INSTIs) or three (control vs. naive vs. INSTIs) groups, as appropriate. Statistically significant p values are in bold. Asterisks indicate statistically significant differences with respect to control group (\*p<0.05, \*\*p<0.01 and \*\*\*p<0.001). AIDS (acquired immunodeficiency syndrome), BMI (body mass index), HS (heterosexual), INSTIs (integrase strand transfer inhibitors-based treatment), MSM (men who have sex with men).

### Reads and contigs distribution

After the sequencing and the quality control, 48.8% of the reads (99,038,911 reads) mapped to viruses (**Fig 1A**) and, of them, 98.8% belonged to phages and 1.2% to eukaryotic virus (**Fig 1C**). Among the rest of the reads, 40% mapped to bacteria, 9.8% to other microorganisms and only 1.4% were unannotated (**Fig 1A**). On the other hand, only 0.4% of the contigs mapped to viruses (**Fig 1B**) and, of them, 88.6% belonged to phages and 11.4% to eukaryotic viruses (**Fig 1D**). Among the rest of the contigs, 93.7% belonged to bacteria, 4.9% to other and 1% to unannotated (**Fig 1B**).

**Fig 1.**
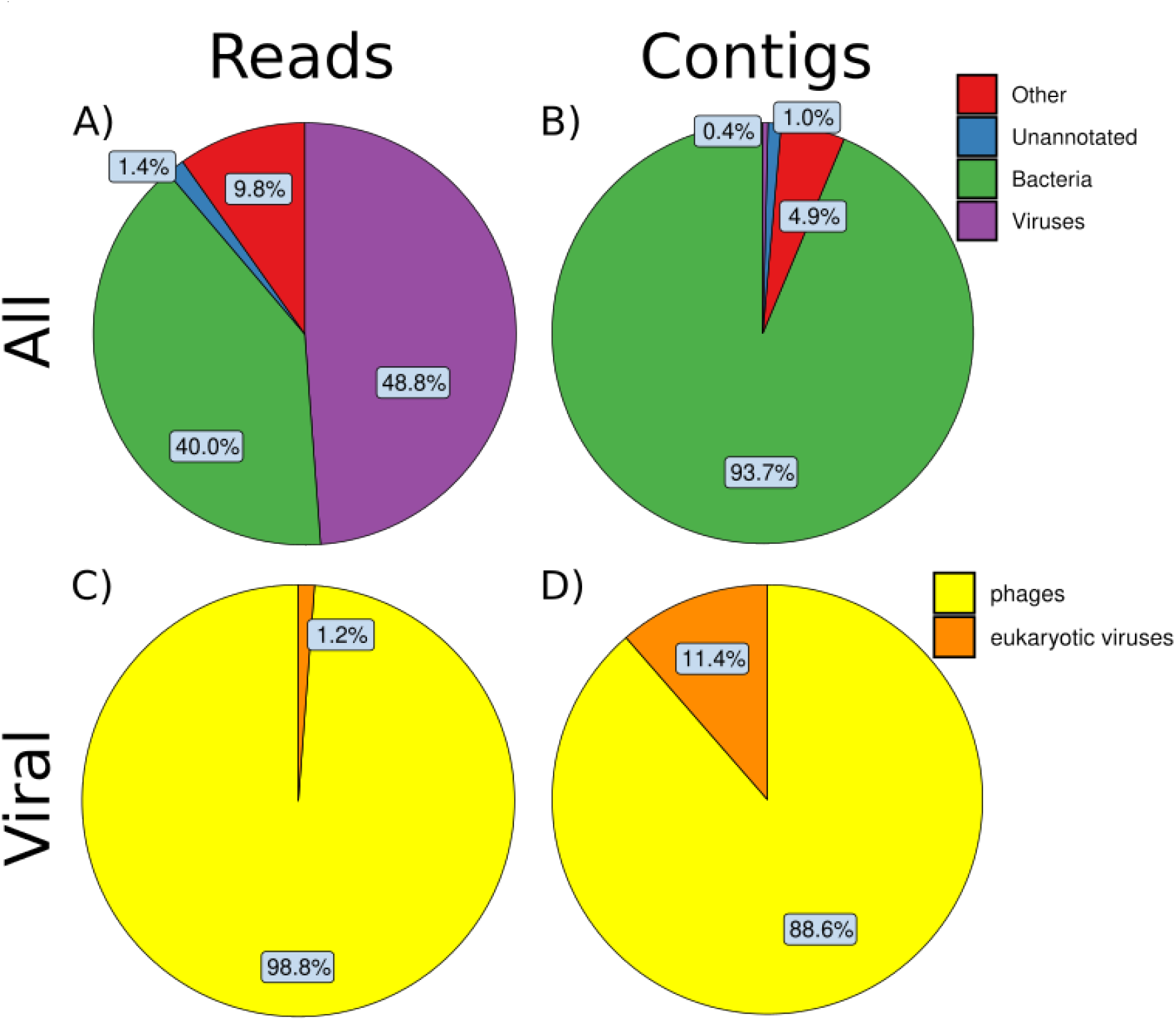
Percentage of the reads (**A** and **C**) and the contigs (**B** and **D**) that correspond to different superkingdoms (**A** and **B**) and viral categories (**C** and **D**).

### Eukaryotic virus diversity and composition

#### 1. Alpha and beta diversity of eukaryotic viruses

The analysis of the α-diversity of eukaryotic viruses did not show statistically significant differences in the indexes analysed (*Observed features*, *Simpson index*, *Shannon index* and *Pielou’s evenness*) (**Fig 2**). However, a slightly decreasing tendence in *Observed features*, *Simpson index* and *Shannon index* in the naive group compared to the controls was observed, which seems to be reverted after INSTIs-based treatment. In the same line as α-diversity, the analysis of β-diversity did not reveal a different clustering between the three groups (**Fig 3**).

**Fig 2.**
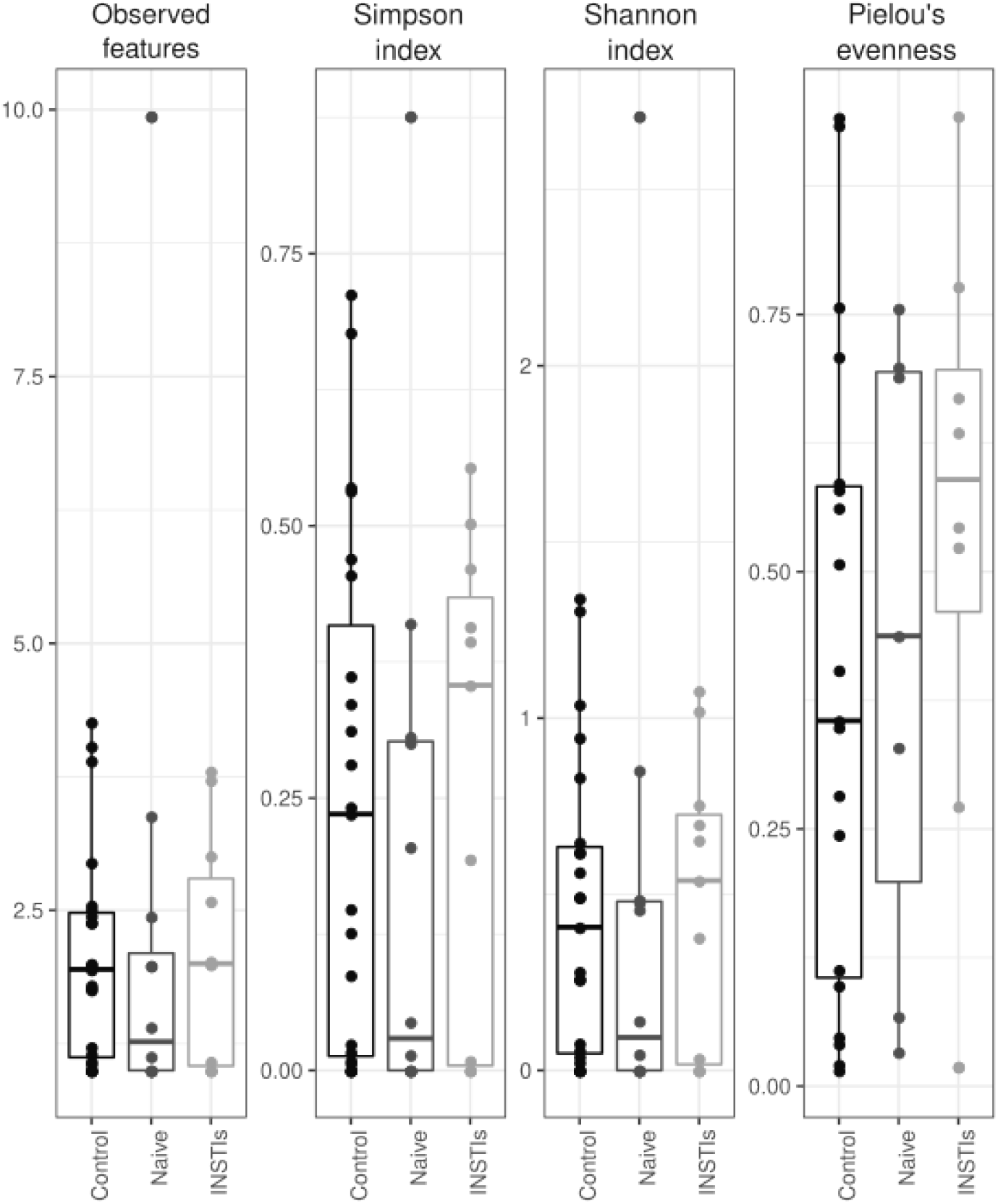
Different indexes of α-diversity from eukaryotic viruses in faecal samples of the studied population. INSTIs (integrase strand transfer inhibitors-based treatment).

**Fig 3.**
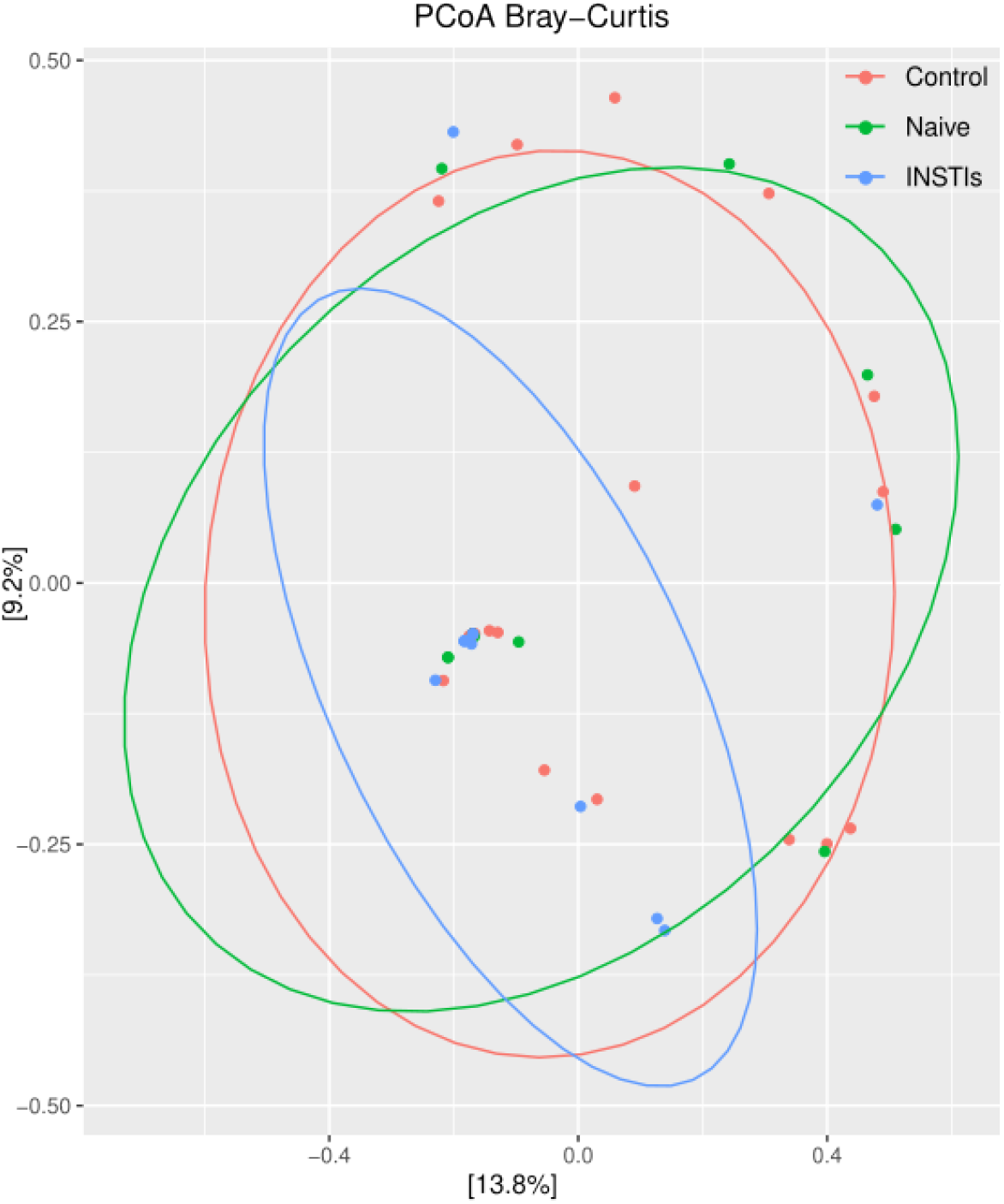
PCoAs from eukaryotic viruses in faecal samples from the studied population (accounting for 23% of the total variation [Component 1 = 13.8% and Component 2 = 9.2%]). Results are plotted according to the first two principal components. Each circle represents a sample: red circles represent the uninfected volunteers, green circles represent the naive group and blue circles represent the INSTIs-treated group. The clustering of sample is represented by their respective 95% confidence interval ellipse. INSTIs (integrase strand transfer inhibitors-based treatment).

#### 2. Distribution of eukaryotic viruses

Eukaryotic viruses were classified into three groups: animal viruses (4 families), plant and fungi viruses (9 families) and small circular viruses (7 families). **Fig 4** shows the presence of these viruses and their relative abundance comparing their presence in controls/uninfected subjects, naive and INSTIs-treated groups to provide a first vision of their distribution. Our data revealed that plant and fungal viruses are the most presented, being those from *Virgaviridae* family the most abundant, followed by small circular viruses. Although plant viruses are likely passengers, they were the only group found in more than 50% of all samples. On the other hand, animal viruses were the least abundant. Due to the low percentage of reads belonging to eukaryotic viruses compared to those belonging to phages and the wide deviation between samples, DESeq analysis did not reveal differentially abundant eukaryotic viruses among the three groups.

**Fig 4.**
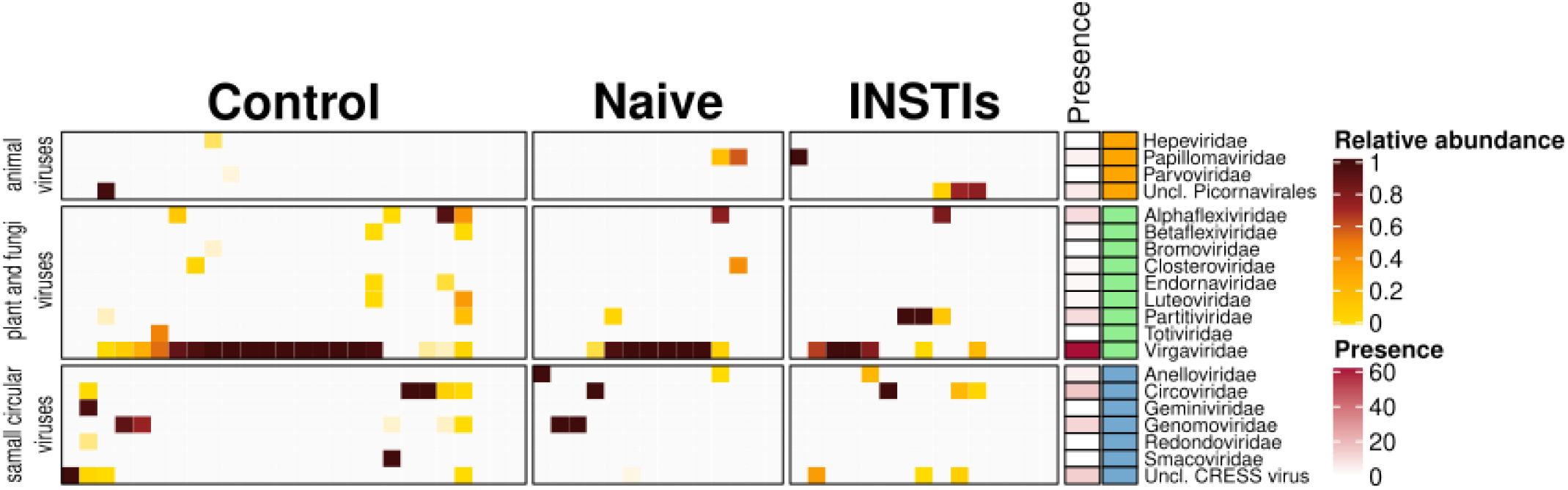
Heatmap of the distribution of eukaryotic viruses between control, naive and INSTIs group. The relative abundance and the presence of the different families between three groups is shown (animal viruses, plant and fungi viruses and small circular viruses). INSTIs (integrase strand transfer inhibitors-based treatment).

### Phage diversity and composition

#### 1. Alpha diversity of phages

A significant decrease in *Observed features* and *Fisher’s alpha* indexes was observed in HIV-naive patients compared to controls (p<0.01 and p<0.05 respectively). Such decrease was not present in INSTIs-treated group when compared to controls (**Fig 5**). No differences were observed either in *Simpson index*, *Shannon index*, or in *Pielou’s evenness*.

**Fig 5.**
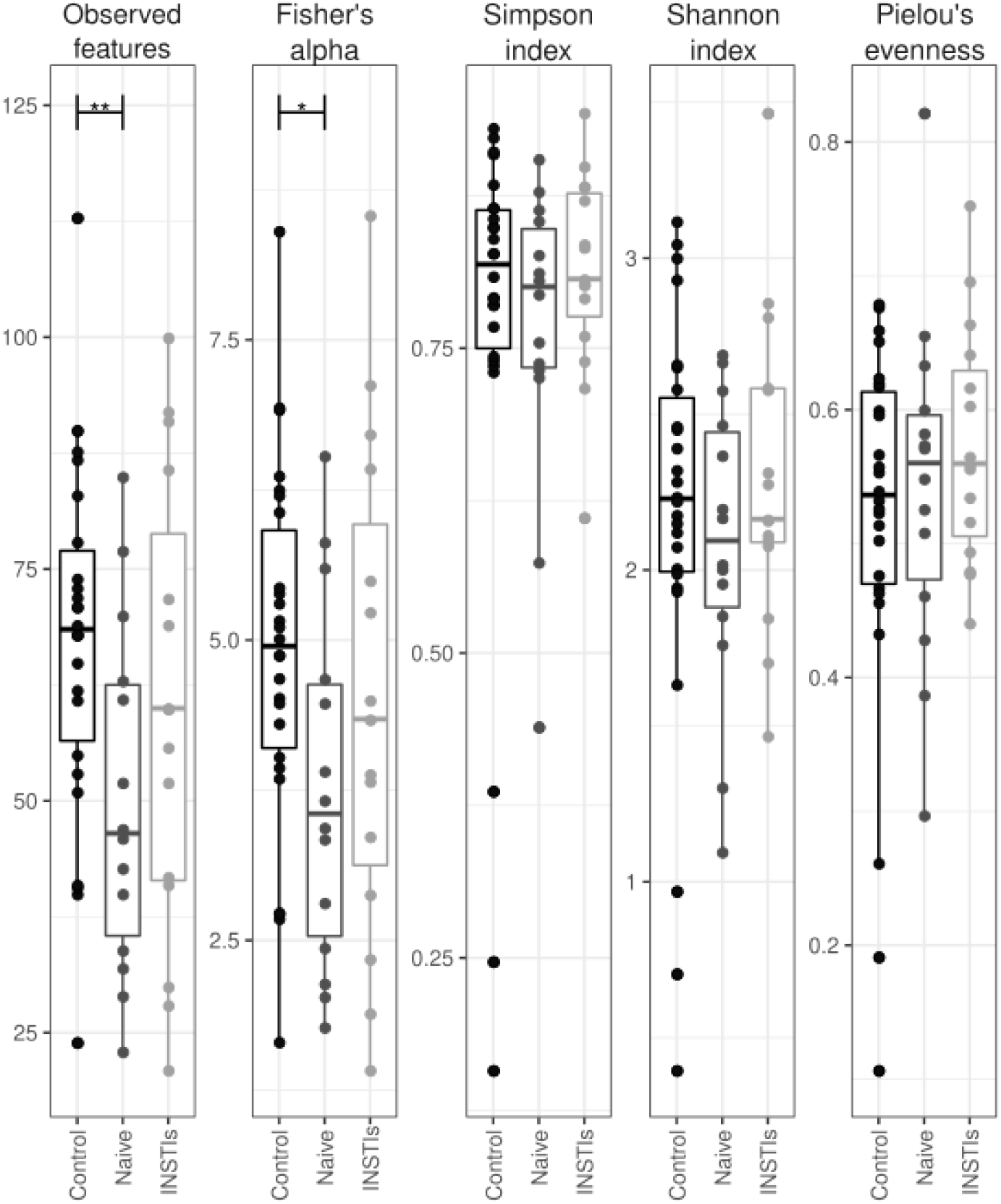
Different indexes of α-diversity from phages in faecal samples of the studied population. *p<0.05 *vs*. control, **p<0.01 *vs*. control. INSTIs (integrase strand transfer inhibitors-based treatment).

#### 2. Beta diversity of phages

**Fig 6** shows the PCoA obtained from the studied population. The control group is clearly different from the naive group (padj<0.01) and the INSTIs-treated group (padj<0.05). However, statistically significant differences were not observed between the naive group and the INSTIs-treated patients in terms of β-diversity, although it is worth mentioning that samples coming from INSTIs-treated patients were more grouped than those coming from naive patients, as can be observed in **Fig 6**.

**Fig 6.**
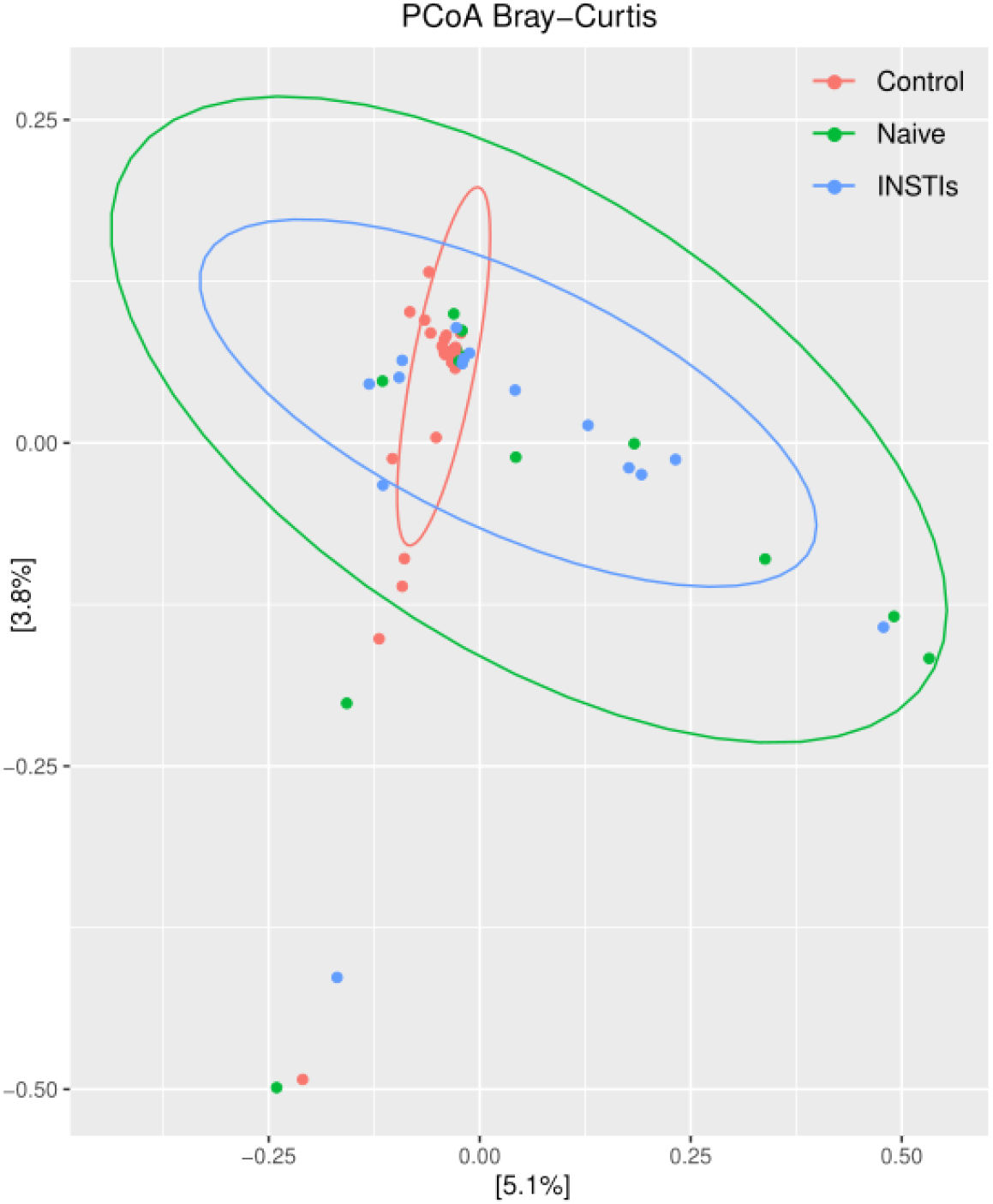
PCoAs from phages in faecal samples from the studied population (accounting for 8.9% of the total variation [Component 1 = 5.1% and Component 2 = 3.8%]). Results are plotted according to the first two principal components. Each circle represents a sample: red circles represent the uninfected volunteers, green circles represent the naive group and blue circles represent the INSTIs-treated group. The clustering of samples is represented by their respective 95% confidence interval ellipse. padj<0.01 naive *vs*. control and padj<0.05 INSTIs *vs*. control. INSTIs (integrase strand transfer inhibitors-based treatment).

#### 3. Differential abundance of phages

DESeq analysis revealed an increase in *Caudoviricetes* class in naive patients compared to the non-infected group and a decrease in *Malgrandaviricetes* class in INSTIs-treated patients compared to the control group. However, differences between naive and INSTIs-treated patients were not detected (**Table 2**).

**Table 2.** Phage taxonomical orders that present a differential abundance in the studied population.

| Control vs. |  |  |  |  |  |  |  |
| --- | --- | --- | --- | --- | --- | --- | --- |
| Naive |  |  |  | INSTIs |  |  |  |
| Category | Taxonomic group |  | padj | Category | Taxonomic group |  | padj |
| Phylum | *Uroviricota* | ↑ | 0.011 |  |  |  |  |
| Class | *Caudoviricetes* | ↑ | 0.011 |  |  |  |  |
|  |  |  |  | Phylum | *Phixviricota* | ↓ | <0.001 |
|  |  |  |  | Class | *Malgrandaviricetes* | ↓ | <0.001 |

#### 4. Lifecycle prediction of phages

Comparison between controls and HIV-infected groups revealed a statistically significant increase in the relative abundance of lysogenic phages in HIV-infected patients (p<0.05 *vs*. control) (**Fig 7A**), that was not reversed after INSTIs-based treatment (**Fig 7B**). These differences couldn’t be solely explained by changes in the relative abundance of *Caudoviricetes* class nor CrAssphages (data not shown).

**Fig 7.**
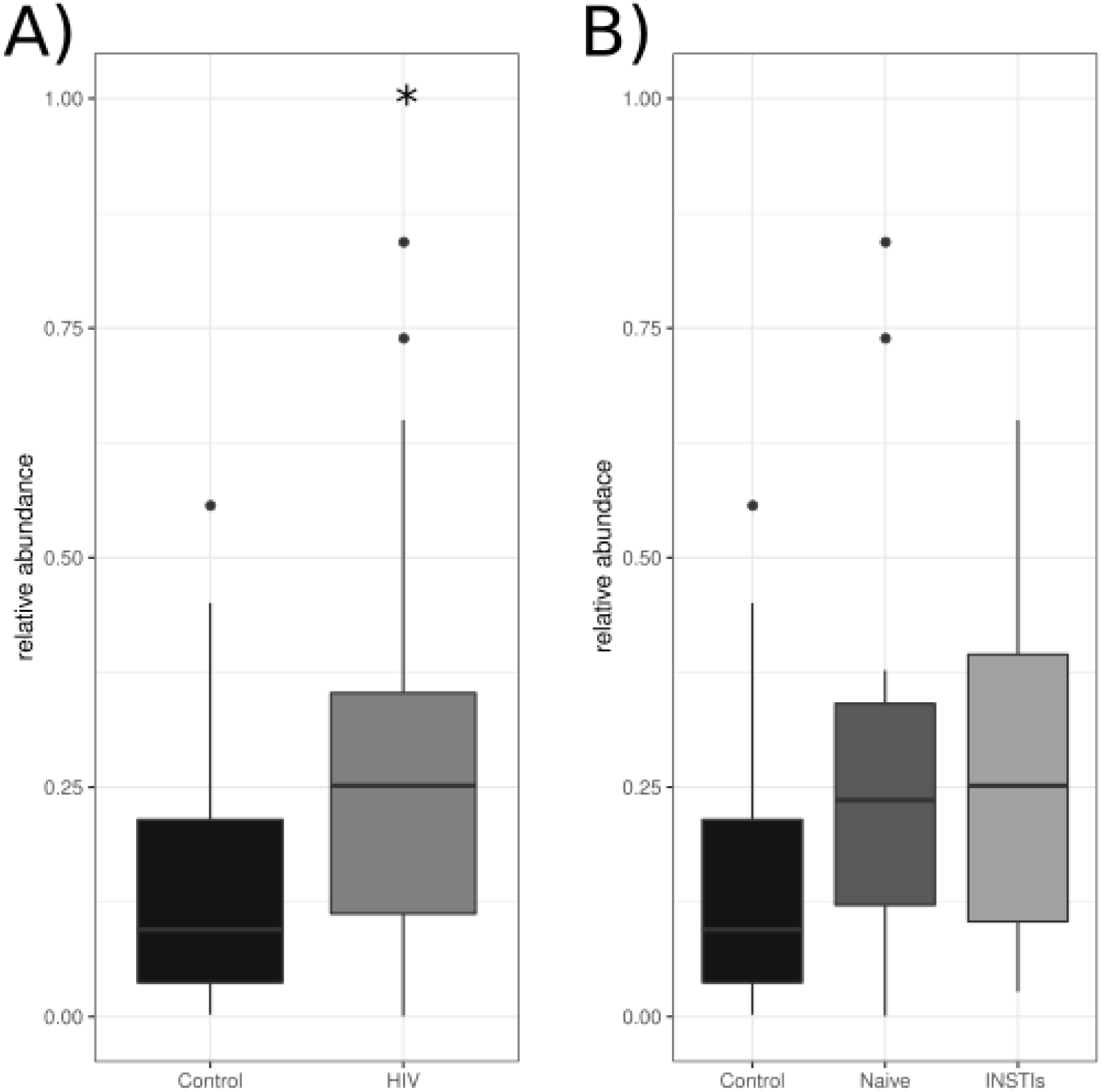
Relative abundance of the lysogenic phages comparing control group *vs*. HIV-infected patients (A) and control group *vs*. naïve patients *vs*. INSTIs-treated patients (B). *p<0.05 *vs*. control. HIV (human immunodeficiency virus), INSTIs (integrase strand transfer inhibitors-based treatment).

#### 5. Host prediction of phages

**Fig 8A** shows the host prediction of the four classes of phages detected (*Caudoviricetes*, *Duplopiviricetes*, *Faserviricetes* and *Malgrandaviricetes*) and of those viruses which were unable to be classified (Unannotated). *Caudoviricetes* were predicted to mainly infect Firmicutes followed by Bacteroidetes phylum. *Malgrandaviricetes* were predicted to mainly infect Proteobacteria phylum followed by Firmicutes and Chloroflexi phyla. The main host of *Duplopiviricetes* and *Faserviricetes* couldn’t be detected and, therefore, was identified as “unknown”.

**Fig 8.**
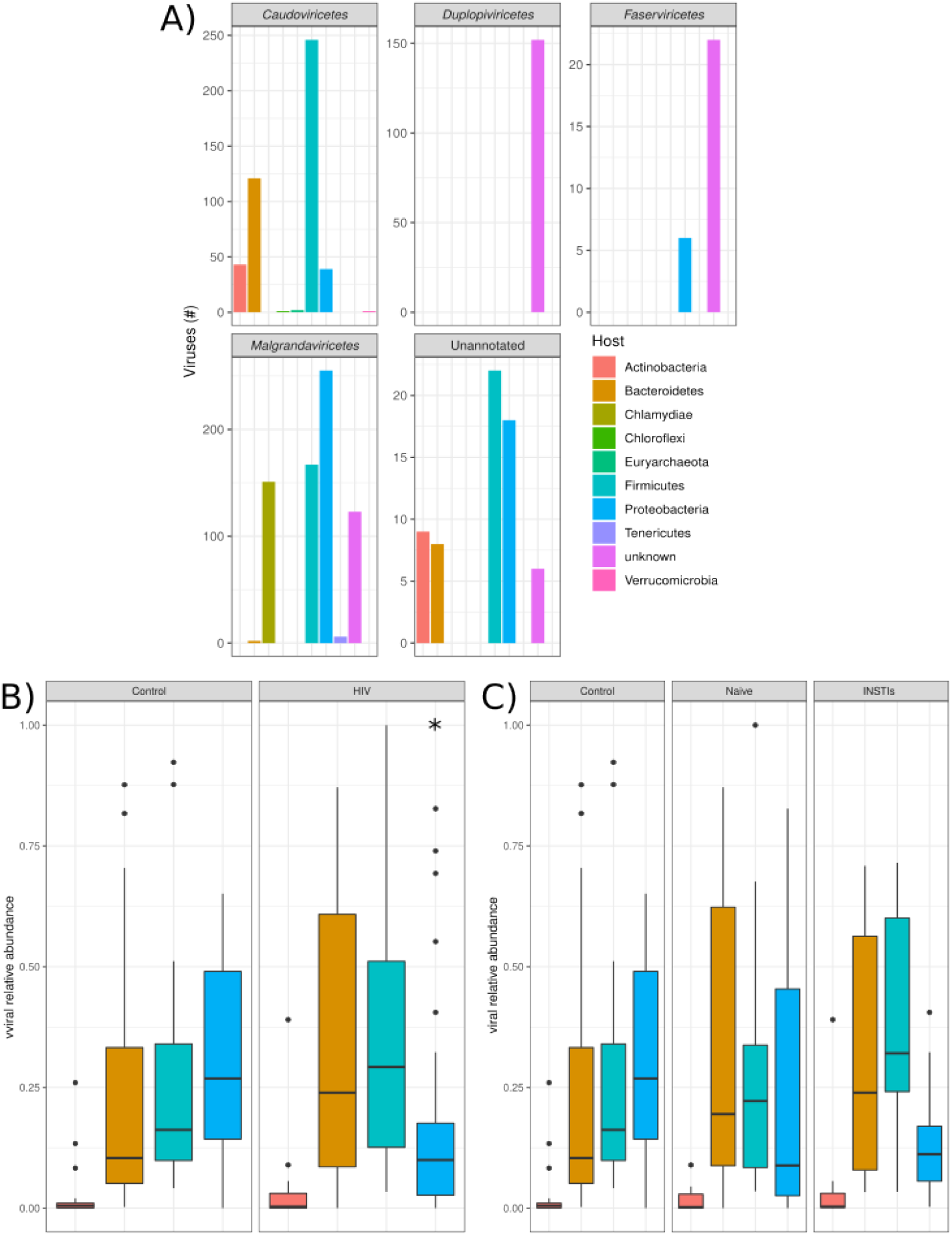
Host prediction of the different phages. **A)** Prediction of which bacteria is infected by phage grouped based on class-level taxonomy. **B)** Differences between control and HIV in the relative abundance of the phages infecting bacterial phyla. **C)** Differences between control, naïve and INSTIs in the relative abundance of the phages infecting bacterial phyla. *p<0.05 *vs*. control. INSTIs (integrase strand transfer inhibitors-based treatment).

Finally, the comparison between the relative abundance of the phages infecting each bacterial phylum revealed that INSTIs-based treatments were not able to restore the decrease observed in the relative abundance of Proteobacteria-infecting phages (p<0.05 HIV *vs*. control) (**Fig 8B and Fig 8C**).

## Discussion

To our knowledge, this is the first time that DNA and RNA viruses have been identified in the gut of HIV-infected people. The impact of INSTIs-based treatments on gut virome composition of HIV-infected people has also been deeply analyzed. This study implies a “snapshot” needed to identify if INSTIs-based treatments are able to restore gut dysbiosis not only at bacterial [13] but also at viral level.

Our study revealed that in our gut samples (stools), most of the viral reads and contigs belonged to phages while eukaryotic viruses were less abundant, which is in line with previous studies that highlight that bacteriophages are the vast majority of the viral component in the human gut [7]. In this context, the lack of differences in alpha and beta diversity of eukaryotic viruses could be due to the low number or reads belonging to these viruses, though a slightly non-significant decrease of richness and diversity indexes in naive patients compared to the controls was observed, along with a partial recovery after INSTIs-based treatments. Besides, our results revealed that plant and fungal viruses (mainly *Virgaviridae*) were the most abundant followed by small circular viruses and animal viruses suggesting that most of the reads belonging to eukaryotic viruses come from diet. These results differ from those obtained by Monaco *et al*. (2016) [10] who reported that *Adenoviridae*, *Anelloviridae, Circoviridae* and *Papillomaviridae* viruses, all of them animal viruses, were the most abundant eukaryotic viruses in the enteric virome of HIV-infected subjects. However, this study only analyzed DNA viruses, so their viral community is quite different from ours, in which we have included both DNA and RNA viruses. Besides, the HIV cohort analyzed in Monaco’s study was different from ours in terms of lifestyle, diet, age, ART and treatment duration (33.27 ± 5.04 months *vs*. 6.7 years).

Regarding bacteriophages, a significant decrease in *Observed features* and *Fisher’s alpha* indexes was observed in naive patients compared to uninfected controls. However, INSTIs-based treatment was able to reverse such decrease, since no significant differences were observed among treated patients and the control group. These results are contrary to those obtained by Monaco *et al*. [10], who reported no significant differences in richness or *Shannon* diversity of bacteriophage families or genera by HIV-infection or treatment status. The differences among both studies could be due to several factors associated with the population recruited and described before and by the fact that only DNA viruses were analyzed in the study of Monaco et al. [10] whereas we have included both DNA and RNA viruses. In fact, phages can be DNA and RNA viruses, therefore, our study is more similar to reality because it assesses the viral community as a whole. Besides, we have revealed a different clustering pattern of HIV-infected patients considering β-diversity that has not been shown before.

Our study has also shown a statistically significant increase in *Caudoviricetes* class in naive patients compared to the control group. *Caudoviricetes* are dsDNA phages that have been revealed to be increased in IBD [17]. Therefore, this increase might be associated to, or be responsible for the increased inflammatory state observed in HIV-infected patients that is abolished after INSTI-based treatment (results from our group submitted for publication). Thus, the role of this viral class deserves more attention in the context of HIV-infection and inflammation. On the other hand, we observed a statistically significant decrease of *Malgrandaviricetes* class in INSTIs-treated patients, although, —up to date— its physiological meaning is unknown. Thus, more studies are needed in order to determine the role of this phage class in gut in the context of HIV-infection and INSTIs-based treatment.

Interestingly, we observed an increase in lysogenic phages related to HIV-infection that was not reversed by INSTIs-based treatments. Our hypothesis that could explain this effect is that inflammation triggered by HIV-infection creates an stressful state in the enteric environment that, by one side reduces bacteriome α-diversity (results from our group submitted for publication); and, on the other side, induces a natural selection in phages towards lysogenic profiles since lytic virions have more difficulties to find a new bacterial host to infect. More studies are needed to confirm such hypothesis. In addition, these effects were not reversed by INSTIs-based treatments, so more studies are required to analyse the possible clinical implications of such findings.

We have also revealed for the first time a significant decrease in Proteobacteria-infecting phages in HIV-infected patients, which was not restored by INSTIs-based regimens. These results are very interesting because they correlate with the increase observed in some Proteobacteria taxonomical orders (specifically Aeromonadales order and *Succinivibrio genus*) observed in naive patients (results from our group submitted for publication). Considering that previous studies suggests that *Succinivibrio* could be associated with defects in gastrointestinal functions, such as diarrhoea and abdominal pain [18–21], a phage therapy focused on decreasing *Succinivibrio* presence could suppose an improvement in quality of life of these patients.

This study certainly has some limitations that will be discussed in the following paragraph. There are some differences between the control group and the naive group, such as gender, age, and smoking habits, all of them factors that could have an impact on GM. However, the two HIV-infected groups were well-balanced taking into account these factors, so the differences observed between them will be independent from these factors and could be attributed to INSTIs-based treatments. Besides, when the control group is compared with all HIV-infected patients (both naive and INSTIs-treated together) no statistically significantly differences were observed in age, so the differences observed in the gut virome among both groups could be attributed to HIV infection. Moreover, we have performed a study comparing the control/uninfected group *vs*. naive group controlled by age (splitting into two groups, “young” and “aged”, according to the median age of the population) and only differences were observed on β-diversity, suggesting that age could be a factor with some kind of impact on gut virome composition. However, since the “aged” group only included 3 naive patients from 19, the potential role of HIV infection more than age *per se* could also contribute to such actions.

To sum up, our study demonstrates that HIV infection has a direct impact on gut bacteriophages and eukaryotic virus abundances, and INSTIs-based treatments were able to reverse the effects observed on α-diversity metrics. These results could serve as a first step for future development of coadjuvant therapies (based not only on probiotics/prebiotics but also phage therapies and/or fecal transplantation) to reduce bacterial dysbiosis in HIV-infected patients and, therefore, to improve the inflammatory state associated with the infection and the long-term consequences of such state. This will undoubtedly improve the health and quality of life of HIV-infected patients.

## Materials and Methods

### Patient recruitment

HIV-infected patients (naive and under ART) were recruited from the Infectious Diseases Department at Hospital Universitario San Pedro (HUSP) (Logroño, Spain) from March 2019 to February 2021. The group of HIV-infected ART-treated patients included HIV-infected patients in first line of treatment to avoid confounding effects due to previous treatments. Treatment was based on INSTIs (dolutegravir or bictegravir) with a backbone based on one or two Nucleoside Reverse Transcriptase Inhibitor (NRTI) for at least one year and with viral load <20 copies/ml in the last six months. All HIV-infected patients were immune responders. The presence of acquired immunodeficiency syndrome (AIDS), via of transmission and coinfection with hepatitis B virus (HBV) and/or hepatitis C virus (HCV) were also registered. In case of coinfection, degree of liver fibrosis was evaluated by FibroScan^®^ (Echosens, Paris, France) method. Patients were classified according to the METAVIR scoring system (F0, no fibrosis; F1, portal fibrosis without septa; F2, portal fibrosis and few septa; F3, numerous septa without cirrhosis; F4, cirrhosis) [22]. CD4+ T-cell, CD8+ T-cell counts, and viral load were measured using flow cytometry (NAVIOS EX, Beckman Coulter) and COBAS 6800 Analyzer (Roche Molecular Systems Inc., Branchburg, New Jersey, USA), respectively, as a clinical procedure in the HUSP. Healthy patients (non-HIV-infected patients) were also recruited as control group (n=26). For both HIV-infected patients and controls, following exclusion criteria were applied: <18 years, patients who do not sign the informed consent, pregnant women, individuals with inflammatory disease in the last 2 months, patients treated with antibiotics, anti-inflammatory drugs, immunosuppressive drugs, statins or probiotics in the last 2 months, individuals with renal insufficiency, patients with neoplasms, individuals with history of intestinal surgery (except from appendectomy or cholecystectomy), IBD, celiac disease, chronic pancreatitis or any other syndrome related to intestinal malabsorption [13]. Patients treated with statins were excluded because it was demonstrated that this therapy can cause gut dysbiosis [23,24]. Finally, weight, height, waist circumference, systolic and diastolic pressure, alcohol consumption, and smoking habits were also registered from all participants. This study was performed following the Helsinki Declaration and was approved by the Committee for Ethics in Drug Research in La Rioja (CEImLAR) (28 February 2019, reference number 349). All participants provided their written informed consent.

### DNA and RNA extraction from stools samples and shotgun sequencing

Fresh stool samples were received at CIBIR, collected, aliquoted in tubes with O-ring caps (45-55mg) and stored at −80°C for further analysis. Later, stool samples were thawed and fecal viral DNA and RNA were extracted using the NetoVIR protocol as described before [25]. The aliquots were suspended in sterile dPBS (10%) and homogenized using the MINILYS homogenizer (Bertin Technologies) for 1 min at 3,000 rpm. Homogenates were centrifuged for 3 min at 17,000g and filtered using a 0.8 μm PES filter (Sartorius). Filtrates were treated with micrococcal nuclease (New England Biolabs) and benzonase (Novagen) at 37°C for 2h. Viral nucleic acids were extracted using the QIAMP^®^ Viral RNA mini kit (Qiagen, Venlo, Netherlands) without addition of carrier RNA to the lysis buffer. Subsequently, random amplification was performed using a modified version of the WTA2 kit (Sigma-Aldrich) with the following parameters: 94°C for 2 min, 17 cycles of 94°C for 30 sec and 70°C for 5 min. The WTA2 products were purified using the MSB Spin PCRapace kit (Stratec Molecular). Quantification of purified product was performed using Qubit™ dsDNA HS Assay kit with the use of a Qubit 2.0 fluorometer (Thermo Fisher Scientific, MA, USA). Sequencing libraries were prepared using the Nextera XT DNA Library kit (Illumina). Sizes of the libraries were checked with the Bioanalyzer 2100 using the High Sensitivity DNA kit (Agilent Technologies, USA).

Sequencing was performed using NextSeq 500 high-throughput Illumina platform (2×150 bp paired-end, Nucleomics Core facility, KU Leuven, Belgium). Computational analysis was performed with ViPER [26]. Ambiguous bases, low-quality reads, primers, and adapter sequences were removed with Trimmomatic (v0.39). Sequences mapping to the “contaminome” were removed using Bowtie2 (v2.4.2) in “very-sensitive” mode [27]. Quality-controlled reads were *de novo* assembled into a set of contigs using MetaSPAdes (v3.15.3) using 21,33,55 and 77 k-mer length [28]. A set of non-redundant scaffolds was obtained by clustering the contigs with a length greater than 500 bp at 95% average nucleotide identity and 85% coverage using CheckV’s clustering scripts [29]. Instead of calculating abundances by mapping the quality-filtered reads to the complete set of non-redundant scaffolds, reads were only mapped against the representatives of the cluster containing a scaffold from that sample to avoid false-positive detection of closely related sequences. Abundances per sample were obtained by mapping the quality-controlled reads back to the set of representative scaffolds using bwa-mem2 (v2.2.1) [30]. Representative scaffolds with a horizontal coverage of 70% or higher were kept for further analyses.

#### 1. Eukaryotic viruses

Eukaryotic viruses were identified and classified by homology-based approaches. The representative scaffolds set was compared against the NCBI nucleotide database using BLASTN (v2.11.0, e-value ≤ 1e-10) [31], and against a non-redundant protein sequence database using DIAMOND (v.2.0.13, sensitive mode) [32] (nonredundant [nr] and nucleotide [nt] databases downloaded from NCBI) and CAT (v5.2.3) [33]. Classification was based on the principle of lowest common ancestor as determined by the ktClassifyBLAST module in KronaTools (v2.8) [34].

#### 2. Prokaryotic viruses

Prokaryotic viruses (bacteriophages) were identified using Virsorter2 (v2.2.3) with score ≥ 0.5 [35]. The completeness of these contigs was determined with CheckV (v0.8.1) [29]. Bacteriophages were selected for further analysis based on a combination of Virsorter2 identification and ≥ 50% completeness. Classification was performed as described in the previous section. Additionally, taxonomic classification was expanded using marker gene approaches as determined by Cenote-Taker2 (v2.1.3) [36]. The lifestyle of bacteriophages was determined based on the appearance of lysogeny-specific genes. These genes were predicted using the functional annotation module of Cenote-Taker2 and can be found in **Supplementary Table 1** [36]. The host of bacteriophages was determined using Random Forest Assignments of Hosts (RaFAH, v0.3). Bacterial hosts were predicted on phylum level with score ≥ 0.14 [37].

### Statistical analysis

Results are presented as mean ± standard error of the mean (SEM) for quantitative variables and as percentage for qualitative variables. Categorical variables were analysed using the Chi-square or Fisher’s exact test. Normal distribution of quantitative variables was checked using the Shapiro-Wilk test. Comparison between two groups were performed using unpaired t test or U-Mann Whitney depending on the normality of the data. Comparison between three or more groups were analysed using ANOVA followed by Tukey post-hoc regardless the normality of the data. P values < 0.05 and false discovery rates (FDRs) < 0.05 were considered as statistically significant. Statistical analysis was performed using GraphPad Prism 8 (GraphPad Prism^®^, La Jolla, California, USA), R software (version 4.0.5) and R Studio (version 1.4.1105).

Alpha and beta diversity were analysed using phyloseq [38]: α-diversity is a measure of sample-level species richness, whereas β-diversity describes inter-subject similarity of microbial composition and facilitates the identification of broad differences between samples. The measure of α-diversity was analysed using *Observed Features*, *Fisher’s alpha*, *Shannon index*, *Simpson index* and *Pielou’s index* with the plot_anova_diversity function of the microbiomeSeq package. *Observed Features* and *Fisher’s alpha* are based in richness, *Pielou’s index* is based in evenness and *Shannon index* and *Simpson index* are based in diversity (richness + evenness). The measure of β-diversity was analysed using Bray Curtis (ordinate function from phyloseq package) and visualized using Principate Coordinate Analysis (PCoA) (plot_ordination function from phyloseq package) and statistically significant differences were evaluated with PERMANOVA (adonis2 and pairwise.adonis functions from the vegan package). Finally, the analysis of the differential composition of microbiomes was carried out with DESeq2 at phylum and class taxonomic levels regarding phages and at phylum, family, and genus taxonomic levels regarding eukaryotic viruses. When required, statistical significance was evaluated with the adequate non-parametric test. All these analyses were performed using R (v3.6.3) and R Studio (v1.2.1335).

## Author contributions

PPM and JAO conceived the study concept. PBN, LM, VI and JA recruited the patients. PVB, MI and ER conducted the prosecution of the samples. PVB, MI and LC extracted the DNA and RNA from stool samples. PVB, JD, LDC and JM carried out the shotgun sequencing and the bioinformatic analysis. PVB, JD, JM and PPM did the statistical analysis. PVB wrote the manuscript. All authors read an approved the final manuscript.

## Compliance with ethics guidelines

This study was performed following the Helsinki Declaration and was approved by the Committee for Ethics in Drug Research in La Rioja (CEImLAR) (28 February 2019, reference number 349). All participants provided their written informed consent.

## Data availability

The datasets generated during and/or analysed during the current study are available in the NCBI SRA repository, http://www.ncbi.nlm.nih.gov/bioproject/819232

## Thanking patient participants

We would like to thank all participants of this study and physicians involved in patient recruitment.

## Supporting information

**Supplementary Table 1.** Predicted lysogenic-specific genes and their functions.

| **Lysogenic-specific genes** | **Function** |
| --- | --- |
| Integrase | Integrase |
| INTEGRASE | Integrase |
| Phage integrase family protein | Integrase |
| SITE SPECIFIC RECOMBINASE XERD | Integrase |
| Excisionase from transposon Tn916 | Excisionase |
| DNA binding domain, excisionase family | Excisionase |
| Repressor protein CI | Repressor |
| P22 AR repressor domain protein | Repressor |
| REPRESSOR PROTEIN | Repressor |
| Cro/C1-type HTH DNA-binding domain | Repressor |
| Phage regulatory protein Rha (Phage pRha) | Repressor |
| Uncharacterized ATPase, putative transposase | Transposase |
| MU TRANSPOSASE | Transposase |
| PROPHAGE LAMBDALM01 ANTIGEN | Prophage domain |
| Mu-like prophage DNA circulation protein | Prophage domain |
| Mu-like prophage I protein | Prophage domain |
| ParB protein | Lysogenic recombination |
| CHROMOSOME PARTITIONING PROTEIN PARB | Lysogenic recombination |

